# Subliminal perception can be predicted from prestimulus activity

**DOI:** 10.1101/2020.01.06.896803

**Authors:** Henry Railo, Roberto Piccin, Karolina M. Lukasik

**Affiliations:** Department of Clinical Neurophysiology, University of Turku, 20521, Turku, Finland; Turku Brain and Mind Centre, University of Turku, Turku, Finland; Department of Psychology, University of Turku, Turku, 20014, Finland; Department of Life Sciences, University of Trieste, 34127 Trieste, Italy; Department of Psychology, Åbo Akademi University, 20500 Turku, Finland

## Abstract

Individuals are able to discriminate visual stimuli they report not consciously seeing. This phenomenon is known as “subliminal perception.” Such capacity is often assumed to be relatively automatic in nature, and rely on stimulus-driven activity in low-level cortical areas. Instead, here we asked to what extent neural activity before stimulus presentation influences subliminal perception. We asked participants to discriminate the location of a briefly presented low-contrast visual stimulus, and then rate how well they saw the stimulus. Consistent with previous studies, participants correctly discriminated with slightly above chance-level accuracy the location of a stimulus they reported not seeing. Signal detection analyses indicated that while subjects categorized their percepts as “unconscious”, their capacity to discriminate these stimuli lay on the same continuum as conscious vision. We show that the accuracy of discriminating the location of a subliminal stimulus could be predicted with relatively high accuracy (AUC = .70) based on lateralized electroencephalographic (EEG) activity before the stimulus, the hemifield where the stimulus was presented, and accuracy of previous trial’s discrimination response. Altogether, our results suggest that rather than being a separate unconscious capacity, subliminal perception is based on similar processes as conscious vison.

## Introduction

One of the major aims of neuroscience is to uncover which neural processes enable a person to consciously experience and respond to stimuli in their environment. Interestingly, visual stimuli that individuals report as not consciously perceived can sometimes influence their behavior, a phenomenon known as subliminal or unconscious perception (Hannula, Simons, & Cohen, 2005; Kouider & Dehaene, 2007; Phillips, 2018; Stein & Peelen, 2021). The neural mechanisms that enable this subliminal capacity are not well understood, and debate continues on whether subliminal perception reveals a truly unconscious capacity that functions independent of conscious perception. In the present study we examine using signal detection theoretic measures to what extent subliminal perception is independent of subjective introspection. Second, we investigate to what extent subliminal perception can be predicted based on the state of the brain before stimulus presentation.

To a large degree, the debate whether subliminal perception is a truly unconscious capacity stems from the difficulty of determining subjective, conscious perception. To measure whether a stimulus was consciously seen, experimenters ask the participant to make a decision, and report what he or she saw. This means that the participant must set some criterion level to which she compares the strength of her subjective perception. When the “signal” (i.e., conscious perception) falls below the criterion for “not seen”, perception is considered unconscious. Subjective perceptual reports can be affected by multiple factors (e.g., decision bias, perceptual bias, motor bias; (Anzulewicz, Hobot, Siedlecka, & Wierzchoń, 2019; Samaha, Iemi, Haegens, & Busch, 2020; Siedlecka et al., 2019)). Different means of reporting conscious perception can yield different results regarding the presence of subliminal perception (Overgaard, Rote, Mouridsen, & Ramsøy, 2006; Ramsøy & Overgaard, 2004), indicating that at least sometimes subliminal perception may only be severely degraded conscious vision (Overgaard, 2011; Phillips, 2020).

According to the currently popular models of vision, the feedforward sweep of stimulus-triggered activation through cortical areas happens outside of the individual’s consciousness, but this activation is assumed to have the power to influence behavior (Vanrullen, 2007; VanRullen & Koch, 2003). The extent to which feedforward processes enable subliminal contents to influence behavior may depend on selective attention: attended stimuli may produce stronger activation, increasing the likelihood of correct behavioral response (Dehaene, Changeux, Naccache, Sackur, & Sergent, 2006). Feedforward activation may also trigger the formation of recurrent activity loops between different cortical areas, enabling stronger subliminal perceptual capacity (Dehaene et al., 2006). These conclusions are supported by studies showing that under certain conditions, where the participants report not seeing a stimulus, early visually evoked activation is preserved but does not ignite widespread cortical activation (Gaillard et al., 2009; Kouider, Dehaene, Jobert, & Le Bihan, 2007; Salti et al., 2015; Supèr, Spekreijse, & Lamme, 2001). Widespread cortical activation is assumed to enable full-fledged conscious perception, allowing the participant to guide behavior based on the conscious contents and, for example, report what they perceived (i.e. “conscious access“; Dehaene & Changeux, 2011; Lamme & Roelfsema, 2000).

If the accuracy of subliminal perceptual responses depends on the strength of the early visual activation (Dehaene et al., 2006), then visually evoked activation should be higher in the trials where the participants provide a correct answer. In line with this prediction, some studies show enhanced event-related potentials (ERP) around 200 ms after stimulus onset in occipitotemporal electrodes when the participants correctly discriminate a target they report not seeing (Koivisto & Grassini, 2016; Lamy, Salti, & Bar-Haim, 2009). However, because reports of conscious vision correlate with ERP amplitude in the same time window—an effect known as visual awareness negativity (Förster, Koivisto, & Revonsuo, 2020; Railo, Koivisto, & Revonsuo, 2011)—this correlate of subliminal perceptions could reflect degraded conscious perception rather than strictly unconscious perception. Koivisto and Grassini (2016) reported that the ERP correlate of subliminal discrimination also correlated with conservative response criterion, suggesting that what the participants labeled “not seen” may have been severely degraded conscious perception. This is consistent with the results of Peters and Lau (2015), which suggest that when perception is measured using a procedure that minimizes the influence of criterion setting, no evidence for strictly subliminal perception is observed (see also, (Rajananda, Zhu, & Peters, 2020)).

Preserved or enhanced early visually evoked neural activation in subliminal trials may not alone explain why in some subliminal trials the participant responds correctly but not in other trials. In contrast to the idea that the strength of early visually evoked responses correlates with the likelihood of correct behavioral responses within reportedly subliminal trials, Salti et al. (2012) found that only the late P3 wave correlated with the accuracy of subliminal discrimination (see also, (Salti et al., 2015)). This could mean that subliminal perception may be less influenced by the strength of the visually evoked response than currently assumed. Possibly the trial-by-trial variation in behavioral performance on subliminal trials is more related to how well decision-making processes can “access” the visual stimulus–related information, rather than how strong early activation the stimulus evokes.

Instead of focusing on visually evoked activity, we reasoned that subliminal perception could be explained by the state of neural activity before stimulus presentation. Neural activity preceding stimulus presentation may influence how well the stimulus is perceived due to multiple factors: The participants’ attentional state may vary spontaneously, but it may also be under voluntary control. Numerous studies have shown that conscious visual perception is strongly influenced by low-frequency oscillations, especially the alpha (8–13 Hz) oscillations, before stimulus presentation (Bareither, Chaumon, Bernasconi, Villringer, & Busch, 2014; Boncompte, Villena-González, Cosmelli, & López, 2016; Britz, Hernàndez, Ro, & Michel, 2014; Iemi & Busch, 2018; Jensen, Bonnefond, & VanRullen, 2012; Kloosterman et al., 2019; Limbach & Corballis, 2016; Mathewson, Gratton, Fabiani, Beck, & Ro, 2009; Samaha, Iemi, & Postle, 2017; Schroeder & Lakatos, 2009; Thut, Nietzel, Brandt, & Pascual-Leone, 2006). However, prestimulus alpha power is currently assumed to influence conscious detection of stimuli, and not perceptual discrimination or subliminal perception (Benwell et al., 2017; Iemi, Chaumon, Crouzet, & Busch, 2017; Samaha et al., 2020).

In the current study, we randomly presented participants with a threshold-level low-contrast stimulus on either the left or right visual field and asked them to report in which visual hemifield the stimulus was presented and then rate how well they saw it (Figure 1A). We reasoned that we would have the highest chance of observing subliminal perception using this type of simple location discrimination task. Moreover, we used lateralized stimuli because we wanted to examine how the pattern of lateralized prestimulus activation influences subliminal perception and evoked visual activity. We used signal detection theoretic measures to test if subliminal perception is independent of subjectively reported visibility.

**Figure 1.**
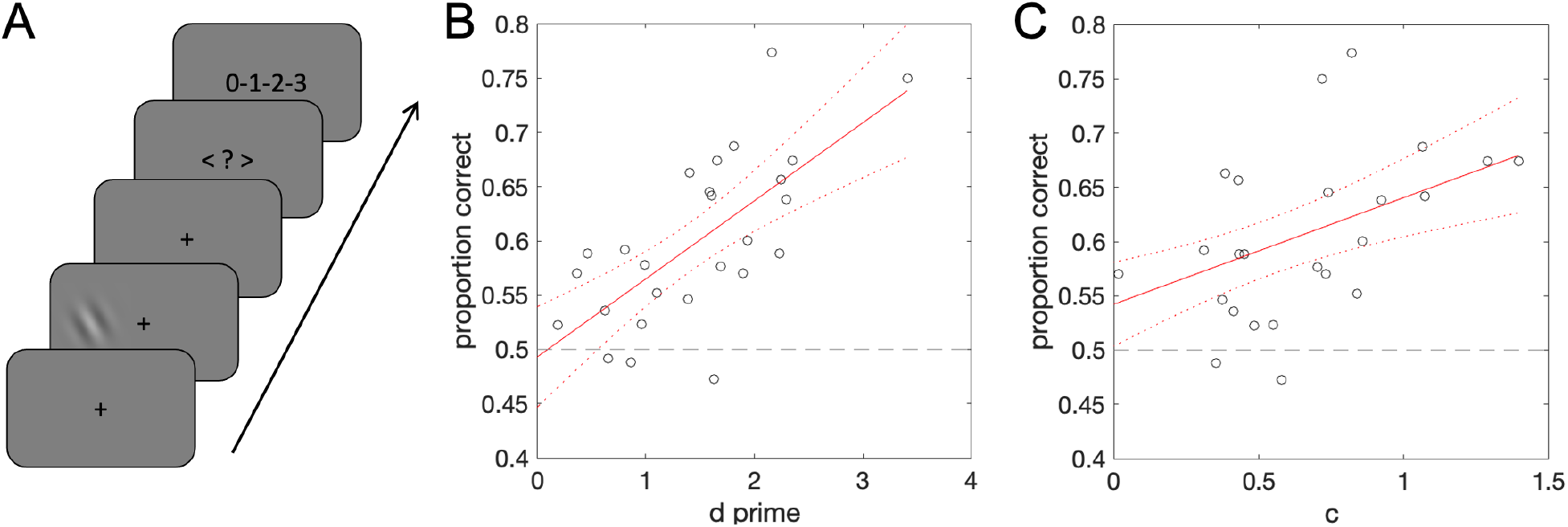
Behavioral paradigm and results. A) A schematic of a single trial: the participants gave a forced discrimination response and rated how well they saw the stimulus. B) Correlation between the sensitivity of subjectively detecting the stimulus and proportion correct (forced discrimination task) on subliminal trials. C) Correlation between criterion for reporting conscious perception and discrimination accuracy on subliminal trials.

## Materials and Methods

### Participants

Thirty-four students (3/34 males, 3/34 left-handed) with a mean age of 24.4 (SD = 3.6) years and no neurological disorders participated in the experiment. Three participants were excluded before the EEG preprocessing because of a high false alarm rate (participants with false alarm rate > 0.3 were excluded; mean false alarm rate of the excluded participants was 0.59, SD = 0.26), and one participant because she always reported seeing the stimuli. The participants were students at the University of Turku. Each participant gave written informed consent. The study was conducted in accordance with the Declaration of Helsinki and was approved by the ethics committee of the Hospital District of Southwest Finland.

### Stimuli

The experiment was run in MATLAB (version R2014b) using the Psychophysics Toolbox (Brainard, 1997). Stimuli were presented on an VIEWPixx/EEG LCD monitor with a 120 Hz refresh rate. The stimuli were Gabor patches (a diameter of 6.5° of visual angle and a frequency of 0.7 cycles/degree; the phase and orientation of the Gabor patch was randomly varied on each trial) presented on a gray 50% background (45.5 cd/m^2^) in left or right hemifield (about 5° from fixation on horizontal meridian). The monitor was not gamma-corrected, which means that average luminance of the Gabor patch does not precisely match the luminance of the background (i.e., stimulus could be detected based on luminance, or contrast information). On two-thirds of the trials, a low-contrast Gabor patch was presented. The intensity of this low-contrast Gabor was individually adjusted to be near a 50% subjective detection threshold using a QUEST staircase (Watson & Pelli, 1983). On one-sixth of the trials, a high-contrast Gabor with three times higher contrast than the contrast of the low-contrast Gabor was presented. The high-contrast stimuli were presented to allow the participant to clearly see the stimuli every now and then, meaning they could use the whole visibility rating scale. Both low-contrast and high-contrast stimuli were presented for a duration of two frames (16.6 ms in total). One-sixth of the trials did not have any stimulus (catch trials), instead containing a blank screen with the fixation point lasting two frames. Only trials with the low-contrast stimuli were included in the statistical analysis.

### Experimental design

The stimulus was presented after a fixation period that varied from 668 ms to 1332 ms from trial to trial to prevent the participant from learning to expect the stimulus at a specific delay. 250 ms after the Gabor was presented (or catch trial), the fixation point turned into an arrow pointing left and right, indicating that the participant should try to report the side of the target. After this, the numbers “0-1-2-3” were presented on the center of the screen to prompt a visibility rating response. Both responses were given by pressing a key on a numpad. The participants were told to try to give their best guess about stimulus location, even when they felt they did not see any stimulus. The visibility rating (Overgaard, 2011; Ramsøy & Overgaard, 2004; Sandberg, Timmermans, Overgaard, & Cleeremans, 2010) was given using a four-step scale where the alternatives were as follows: 0) “did not see stimulus at all,” 1) “not sure but possibly saw something,” 2) “pretty sure I saw it,” and 3) “saw the stimulus clearly.” The difference between alternatives 0 and 1 was stressed to the participant. That is, we emphasized that if they felt that they saw any glimpse of the stimulus, even an extremely weak one, they should select the second lowest alternative. The rating task was preferred over a yes/no task to make sure that the participants used a strict criterion for reporting no subjective perception of stimuli. Throughout the present study, a participant is assumed to be subliminal of a stimulus only when she/he chose the lowest visibility rating (the three higher alternatives are all counted as conscious). No instruction was given about the response speed (of either response). The participants were also told that in some trials, the stimulus would not be presented, and they were told that on these trials, the correct response was to report that they did not see the stimulus. The experiment comprised a total of 400 trials per participant, subdivided into 10 blocks of 40 trials each.

### EEG recording and preprocessing

A 64-channel EEG was recorded at a 500 Hz sampling rate with a NeurOne Tesla amplifier. Before the recording, electrode impedances were brought near 5 kΩ. Two additional electrodes were used to record electro-oculograms (EOG): one electrode was placed beneath and the other beside the participant’s left eye.

Preprocessing was carried out with MATLAB (version R2016b) using the EEGlab toolbox (version 14.1.1b). For each participant, bad channels (channels with no response or very noisy signals) were manually checked and interpolated. EEG data were re-referenced to the average of all electrodes. A 0.25 Hz high-pass filter and an 80 Hz low-pass filter were applied to the data. Line noise was cleaned up using the CleanLine EEGlab plugin. Data were epoched from −2000 ms to 1500 ms relative to the stimulus onset. Trials with eye movements (EOG electrodes) within −500 ms to 500 ms relative to the stimulus onset were discarded (on average, 24 trials, SD = 37 trials were rejected per participant). The locations of the electrodes were transposed so that the left hemisphere was always contralateral and the right hemisphere always ipsilateral relative to stimulus presentation (we present non-transposed results in supplementary figures). A time frequency analysis was performed on single trials using complex Morlet wavelets (frequency range: 1–30 Hz with 2 Hz step size, with the number of cycles increasing linearly from 1 to 12). No baseline correction was applied when examining prestimulus oscillations.

Due to the limited temporal resolution of the wavelet analysis, post-stimulus time windows could influence the prestimulus data points, especially for low (< 3 Hz) frequencies. To control for this, we also calculated power spectral densities from prestimulus (single trial) data segments. The EEG power spectrum reflects not only periodic oscillatory signals, but also aperiodic activity with 1/f-like distribution. We estimated exponent and offset of aperiodic power across frequencies 1–40 Hz using fooof package (Donoghue et al., 2020).

### Statistical analysis

Behavioral signal detection measures of sensitivity (d’) and criterion (c) were calculated from binarized subjective visibility ratings. A hit was defined as a visibility rating of 1–3 when a stimulus was presented. False alarms were trials where the participant reported seeing a target (rating 1–3) when no stimulus was presented.

Influence of prestimulus EEG power on behavior was examined as follows. First, time points between 1000–0 ms before stimulus onset in steps of 24 ms were statistically analyzed to examine if prestimulus oscillatory power predicted behavioral performance. To see which electrode clusters and frequencies may enable predicting subliminal perception, these analyses were performed in a mass univariate manner (Maris & Oostenveld, 2007), analyzing all the time points and all frequencies between 1 and 30 Hz. Statistical analyses were performed on the single-trial data. We used mixed-effects logit models to test whether log transformed oscillatory power at specific time point and frequency predicted the accuracy of the response (correct/incorrect) or subjective visibility (saw/did not see). The mixed-effects models included random intercepts for participants. More complicated models often failed to converge, and the models also took very long to compute. Log transformed power was used as the predictor because untransformed data were strongly right skewed.

Statistical significance was assessed using permutation testing by randomly shuffling condition labels (1000 iterations). We used threshold free cluster enhancement (TFCE) to take into account the clustering of effects in neighboring time points and frequency bands (Smith & Nichols, 2009). Maps of the t values obtained from the mass univariate analyses were TFCE transformed using a function from the Limo EEG package (Pernet, Chauveau, Gaspar, & Rousselet, 2011), using default parameters (E = 0.5, H = 2, dh = 0.1). TFCE was performed for the authentic data and also after each permutation. The highest TFCE score (TFCE_max_) of each permutation was saved to obtain a null distribution corrected for multiple comparisons. This analysis was run separately on data from five clusters of electrodes. To ensure two-tailed testing, and take into account that analysis was repeated five times (for each cluster of channels), we further used a Bonferroni corrected alpha level to assess statistical significance. TFCE values with the probability < .005 when compared with the TFCE_max_ distribution were considered statistically significant.

We also analyzed ERPs and event-related spectral perturbations (ERSP) to examine how behavioral performance (subjective and objective measures) was associated with stimulus-evoked EEG responses. These analyses were performed using a similar mass univariate approach as described earlier (mixed-effects logit models with random intercepts, 1000 permutations, and TFCE, and p = .005 alpha level). In the ERP analyses TFCE was calculated over neighboring time samples and electrodes. Before ERP analysis, EEG was low-pass filtered at 40 Hz. Single-trial ERPs were baseline corrected to −200 ms. In standard ERSP analysis, multiple trials are averaged, and it is measured how EEG oscillatory power changes in response to a stimulus relative to some baseline state. For each trial, we subtracted from each post-stimulus time point and frequency the average prestimulus power (at the corresponding frequency) between −1000 and −100 ms. To reduce computation time, the ERSP was resampled to 24 ms steps for the statistical analysis.

Number of trials per participant in the statistical analyses are as follows. Participants had on average 125 trials in the conscious condition (SD = 42 trials), and 120 (SD = 39) trials in the subliminal condition. When both seen and unseen trial are included in the analysis, the comparison of correct vs. incorrect objective discrimination is based on 190 (SD = 37) vs. 55 (SD = 24) trials per participant on average, respectively. The main analysis of objective discrimination performance on subliminal trials is based on average on 71 (correct response, SD = 23) vs. 49 (incorrect response, SD = 20) trials per participant.

Datasets are available for download at https://osf.io/xz8jr/.

## Results

### Behavioral results

Three participants displayed a very strong rightward response bias: When they reported not seeing a stimulus, they always pressed that the stimulus was presented on the right side. Because the aim of this study was to see if participants who report not seeing a stimulus correctly *guess* the location of the stimulus, these participants were excluded from the behavioral and EEG analysis. The remaining participants did not, as a group, show systematic lateral biases: When no stimulus was presented, these participants responded that the stimulus was on the right hemifield on average on 52% of trials (SD = 2%; one sample t test against 50% level: t = 0.74, df = 26, p = 0.46). Visibility ratings of left and right hemifield targets did not differ statistically significantly (t = 1.13, df = 26, p = 0.26). That these analyses are based on aggregated data, not single trials as the EEG results presented below.

On average, the participants reported seeing the target (visibility rating 1–3) in 57% of the trials (SEM = 3%) when the stimulus was presented. When the stimulus was not presented, they rarely reported seeing the target (mean = 19%, SEM = 4%). When the participants reported not seeing the target at all (visibility rating = 0), they nevertheless correctly reported the side of the stimulus in 59% of the trials on average (t test against 50% chance-level: t = 6.14, df = 26, p = 1.71 × 10^−6^)—we refer to this as “subliminal perception”.

Do accurate responses in the trials where the participants report no conscious perception reveal the existence of a perceptual capacity that functions independently of conscious vision? We calculated the participants’ sensitivity and criterion for *consciously* detecting the target (i.e., the measures were calculated from subjective reports, not objective performance, using signal detection analyses). As shown in Figure 1B, the participants who were not sensitive to subjectively detecting the stimulus (d’ ≈ 0) did not reveal subliminal perception (intercept of the linear model = .49; effect of d’, Beta = 0.072, t = 5.17). As shown in Figure 1C, the participants also tended to use a conservative criterion for reporting conscious perception. This is common with low-contrast stimuli because the participants try to minimize false alarms. Based on extrapolating the fitted linear model, if the participants used statistically optimal criterion (c = 0) for reporting consciousness, they would also have performed at near the chance level in discriminating the stimulus side (intercept = .54; effect of criterion, Beta = 0.098, t = 3.76). Altogether, these results suggest that subliminal perception was not separate from subjectively reported vision.

### Prestimulus EEG correlates

We tested if prestimulus EEG power predicted the participants’ objective discrimination performance when they subjectively reported not seeing the target at all (the lowest visibility rating). Because the lateralization of cortical excitability varies depending on whether attention is focused toward the right or left visual hemifield (Boncompte et al., 2016; Thut et al., 2006; Worden, Foxe, Wang, & Simpson, 2000), we took this into account by transposing the electrode locations for each trial so that the left electrode locations represent electrodes that were contralateral with respect to the visual stimulus (and right electrodes were ipsilateral). The electrode clusters are visualized in Fig. 2A.

**Figure 2.**
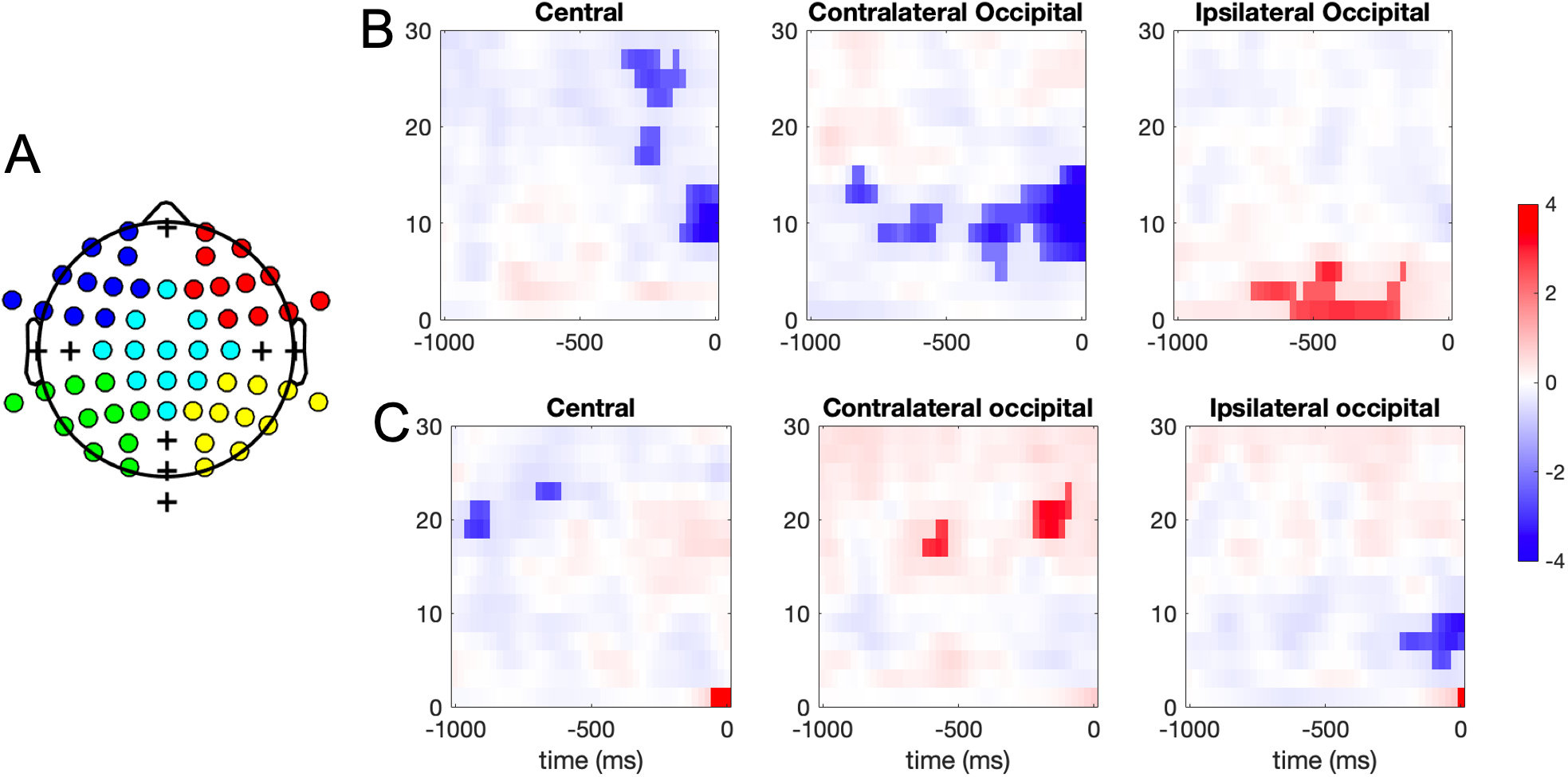
How prestimulus oscillatory power predicts behavior. A) Electrode clusters used in EEG analysis. Each color represents one cluster. Plus-symbols are electrodes that are not part of any cluster. B) Results of mass univariate mixed-effects logit models where a single trial’s prestimulus EEG power was used to predict the location discrimination accuracy on subliminal trials (visibility = 0) and C) binarized visibility ratings (did not see = 0; 1–3 = saw). Color is the t value of the mixed-effects logit model. The blue color denotes a negative correlation and red a positive correlation. The faint colors are time points/frequencies where the model does not statistically significantly predict behavior. Bright colors show statistically significant modulation according to TFCE and permutation testing. Frontal channel clusters are not shown because no statistically significant effects were observed.

As shown in Figure 2B, objective performance on trials where the participants reported not seeing the stimulus at all could be predicted from prestimulus power. Oscillatory power at the occipital electrode clusters predicted the accuracy of responses almost up to 1 s before the stimulus was presented, even though the participants reported just guessing. The effect is lateralized: oscillatory power 8–12 Hz contralateral to the stimulus and 1–5 Hz in ipsilateral occipital electrodes predicts better performance. Statistically significant effects are observed in the occipital electrode clusters, but the frontal and central electrode clusters showed a similar pattern of effects. When electrode locations were not transposed, a weaker prestimulus effect in the alpha band in occipital and central electrodes (up to 250 ms before stimulus onset) on unconscious discrimination was observed (Supplementary Figure 1).

The prestimulus power in single trials was also associated with reported conscious visibility of the target in a trial-by-trial manner (Fig. 2C), but the effect was clearly smaller when compared to effects on unconscious discrimination. A similar effect is found when non-transposed electrode locations were used (Supplementary Figure 2).

To rule out the possibility that the observed prestimulus correlates (Fig. 2B) were caused by temporal smearing of the wavelet, we calculated power spectral density from single trial prestimulus data segments (−500 to −24 ms). We then ran a mixed-effects logit model where we predicted subliminal discrimination accuracy in single trials based on the average power spectral density of the ipsilateral electrode cluster (1–3 Hz), and the average spectral density in the contralateral electrode cluster (8–12 Hz). Oscillatory power in contralateral occipital electrode clusters predicted subliminal discrimination performance, replicating previous analyses (t = −3.05). Ipsilateral occipital power did not predict subliminal discrimination accuracy (t = 1.30). The association between contralateral/ipsilateral occipital power spectral density and subliminal perception are visualized in Figure 3A and 3B.

**Figure 3.**
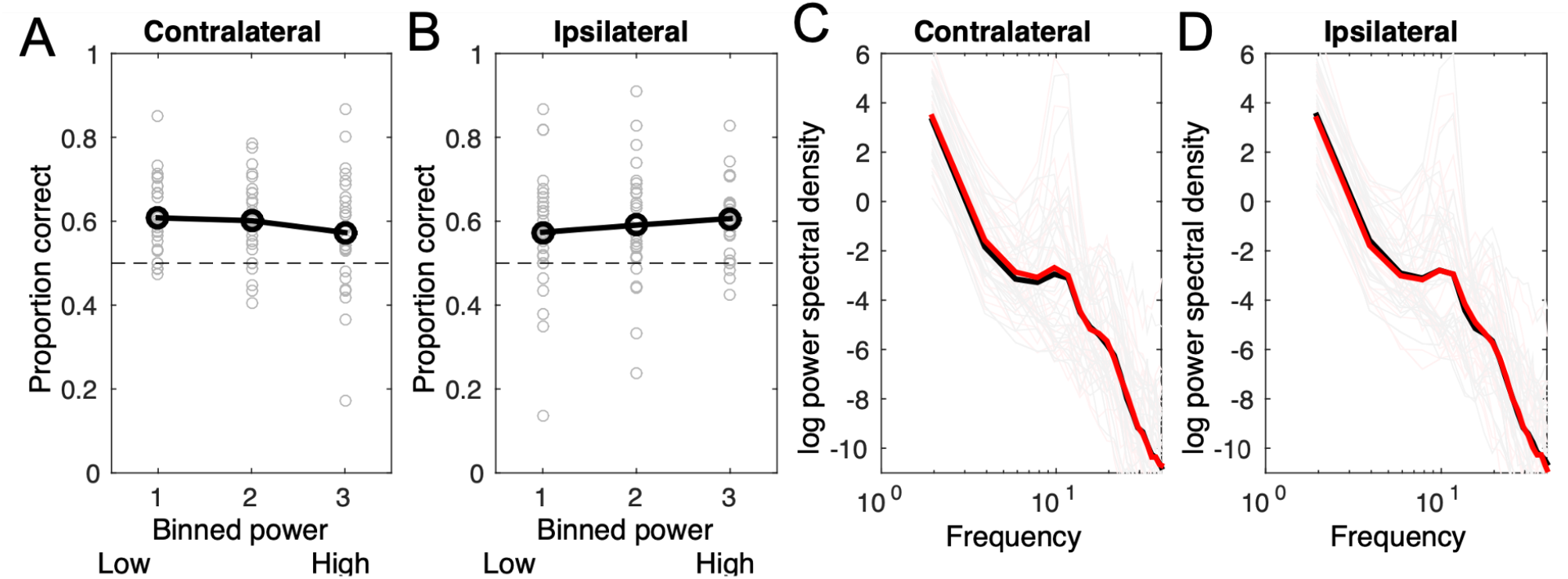
Influence of contralateral and ipsilateral occipital power on subliminal discrimination performance. Panel A) shows the contralateral electrode cluster, and panel B) the ipsilateral electrode cluster. Line and strong data points show the average across participants, and faint symbols individual participants’ data (average across trials). Panels C) and D) show the average power spectrum in contralateral and ipsilateral electrode clusters, respectively. Black line is the condition where the location of a subliminal target was correctly discriminated. Red is incorrect discrimination on subliminal trials.

Next, we examined how well oscillatory power at these two frequencies/electrode clusters predict subliminal discrimination accuracy when confounding variables are controlled for. First, we included in the model the hemifield in which the stimulus was presented to control for the possibility that the effect was caused by a spatial bias. Interaction between stimulus hemifield and contralateral or ipsilateral power are not included in the model presented below because these did not improve the model. Second, accuracy of previous trial’s (whether subliminal or conscious) response was included in the model to account for temporal correlations throughout the experiment (see, e.g., Schaworonkow, Blythe, Kegeles, Curio, & Nikulin, 2015). Predictors related to trial number, participant’s response in the previous trial (left/right), and the stimulus hemifield in the previous trial were initially included in the model, but left out because they failed to explain variation in subliminal discrimination. Finally, to this model we added stimulus hemifield, contralateral power and ipsilateral power as participant-wise random-effect predictors to take into account individual variation in these factors. As shown in Table 1, subliminal discrimination was more accurate in the right hemifield (t = 2.06). The participant was more likely to correctly discriminate the subliminal stimulus when he/she responded correctly in the previous trial (t = 3.89). Even when these factors were included in the model, contralateral occipital prestimulus power predicted subliminal discrimination accuracy (t = − 2.35). Ipsilateral power did not statistically significantly predict subliminal discrimination accuracy in this model (t = 1.21). The area-under-the-curve (AUC) of this model was .70. The same model with shuffled labels produced a maximum AUC of .60 (out of 1000 permutations), and a model with only the intercept term had an AUC of .58.

**Table 1.** Results of a mixed-effects logit model predicting subliminal discrimination accuracy (df = 3291)

| Name | Estimate | SE | t | p | CI (lower) | CI (upper) |
| --- | --- | --- | --- | --- | --- | --- |
| Intercept | -0.322 | 0.214 | -1.500 | 0.134 | -0.742 | 0.099 |
| Contralateral power | -0.043 | 0.018 | -2.359 | 0.018 | -0.078 | -0.007 |
| Ipsilateral power | 0.021 | 0.017 | 1.217 | 0.224 | -0.013 | 0.054 |
| Stimulus in right hemifield | 0.833 | 0.404 | 2.060 | 0.039 | 0.040 | 1.626 |
| Accuracy in previous trial | 0.342 | 0.088 | 3.895 | < .001 | 0.170 | 0.515 |

The above results suggest that behavioral performance in subliminal trials correlates with prestimulus power in specific low-frequency bands. To examine whether these effects stem from true oscillatory activity, or changes in broadband aperiodic activity (Donoghue et al., 2020), we examined to what extent markers of aperiodic EEG activity predict subliminal discrimination accuracy. The aperiodic components (exponent and offset) were calculated separately for occipital ipsilateral and contralateral electrode clusters (from prestimulus data segments). Table 2 shows to what extent the markers of aperiodic activity predicted subliminal discrimination. Neither contralateral alpha (t = −1.34) or ipsilateral delta (t = −0.30) oscillatory power predicted subliminal discrimination accuracy when they were included in the model together with the other predictors. In addition to this “full model” (BIC = 13652), a pruned model (BIC = 13636) that does not include these two predictors is presented in Table 2. In the pruned model, only the offset of estimated aperiodic spectrum predicted subliminal discrimination: weaker contralateral (t = −2.11) and stronger ipsilateral (t = 1.87) broadband activity predicted subliminal discrimination. This finding is important because it suggests that the prestimulus EEG feature that predicts subliminal discrimination may not be an oscillation, but the strength of non-oscillatory broadband EEG.

**Table 2.** Results of mixed-effects logit models predicting subliminal discrimination accuracy (df = 3182)

| Full model |  |  |  |  |  |  |
| --- | --- | --- | --- | --- | --- | --- |
| Name | Estimate | SE | t | p | CI (lower) | CI (upper) |
| Intercept | 0.357 | 0.221 | 1.613 | 0.107 | -0.077 | 0.790 |
| Contralateral exponent | 0.012 | 0.067 | 0.172 | 0.863 | -0.120 | 0.143 |
| Ipsilateral exponent | -0.053 | 0.064 | -0.839 | 0.402 | -0.178 | 0.071 |
| Contralateral offset | -0.068 | 0.059 | -1.162 | 0.245 | -0.184 | 0.047 |
| Ipsilateral offset | 0.110 | 0.060 | 1.832 | 0.067 | -0.008 | 0.228 |
| Contralateral power | -0.025 | 0.019 | -1.341 | 0.180 | -0.062 | 0.012 |
| Ipsilateral power | -0.006 | 0.018 | -0.307 | 0.759 | -0.041 | 0.030 |
| Pruned model |  |  |  |  |  |  |
| Name | Estimate | SE | t | p | CI (lower) | CI (upper) |
| Intercept | 0.340 | 0.224 | 1.518 | 0.129 | -0.099 | 0.779 |
| Contralateral exponent | 0.051 | 0.060 | 0.849 | 0.396 | -0.067 | 0.169 |
| Ipsilateral exponent | -0.041 | 0.059 | -0.693 | 0.488 | -0.155 | 0.074 |
| Contralateral offset | -0.108 | 0.051 | -2.118 | 0.034 | -0.209 | -0.008 |
| Ipsilateral offset | 0.093 | 0.050 | 1.877 | 0.061 | -0.004 | 0.191 |

We also examined if occipital prestimulus power spectral density data predicted reported stimulus visibility. First, we tested if contralateral power and ipsilateral power (same frequencies and time-windows as previously) predicted stimulus visibility when confounding variables are included in the model. The results of the model (which included stimulus hemifield, contralateral power and ipsilateral power as participant-wise random-effect predictors) are presented in Table 3. Here, ipsilateral prestimulus power was negatively associated with reported stimulus visibility (t = −2.25), and again, correct discrimination of the target in previous trial predicted higher likelihood of conscious detection (t = 3.08). Contralateral power did not predict conscious perception (t = −0.49). This is noteworthy because in the comparable model concerning subliminal discrimination (Table 1), contralateral (but not ipsilateral) occipital prestimulus power predicted better performance. This suggests that different prestimulus neural processes may mediate subliminal discrimination and subjective visibility (although based on our behavioral results, they both lie on the same continuum of conscious vision).

**Table 3.** Results of a mixed-effects logit model predicting stimulus visibility (df = 6846)

|  |  |  |  |  |  |  |
| --- | --- | --- | --- | --- | --- | --- |
| Name | Estimate | SE | t | p | CI (lower) | CI (upper) |
| Intercept | -0.204 | 0.132 | -1.548 | 0.122 | -0.462 | 0.054 |
| Contralateral power | -0.008 | 0.016 | -0.493 | 0.622 | -0.039 | 0.023 |
| Ipsilateral power | -0.030 | 0.013 | -2.250 | 0.024 | -0.055 | -0.004 |
| Accuracy in previous trial | 0.192 | 0.062 | 3.080 | 0.002 | 0.070 | 0.315 |
| Stimulus in right hemifield | 0.032 | 0.081 | 0.392 | 0.695 | -0.127 | 0.191 |

Aperiodic prestimulus activity did not predict stimulus visibility, and hence the results are not presented further (in a model with four aperiodic predictors, highest absolute t value = −0.6).

### Event-related potentials

What is the mechanism through which prestimulus power influences the discrimination of subliminal stimuli? High-amplitude low-frequency oscillations have been associated with suppressing the processing of nonrelevant signals, whereas low-amplitude low-frequency power is associated with amplifying sensory input (Schroeder & Lakatos, 2009). The lateralized contribution of occipital prestimulus power on subliminal discrimination performance suggests that successful behavioral responding in subliminal trials may be related to enabling strong stimulus-evoked responses to the (contralateral) visual stimulus while suppressing unwanted signals in the opposite (ipsilateral) visual field (where no target is presented). If this is the case, we should observe enhanced visually evoked responses to correctly discriminated subliminal targets (when compared with the trials of incorrect responses).

To examine how subliminal discrimination accuracy and subjective visibility of the stimulus influences stimulus-evoked activity, we analyzed ERPs in a single-trial manner. As in the prestimulus analyses, the electrode locations in these analyses were transposed so that in the scalp maps, the left hemisphere locations correspond to the contralateral hemifield. As shown in Figure 4A, subjective visibility of the stimulus was associated with a strong increase of ERP amplitudes 275 ms after stimulus onset, spanning most electrode locations. Nearly identical effects are produced by when objective discrimination performance is examined (all trials included, Supplementary Figure 3), in line with the behavioral result that objective discrimination was not independent of subjectively reported vision. However, as shown in Fig. 4B, no strong, statistically significant effects are seen in ERPs when the analysis is restricted to subliminal trials. Note however, that Figure 4B suggests that ERP amplitudes are more negative in contralateral occipital electrodes around 175–275 ms, although this effect does not reach statistical significance in mass univariate testing.

**Figure 4.**
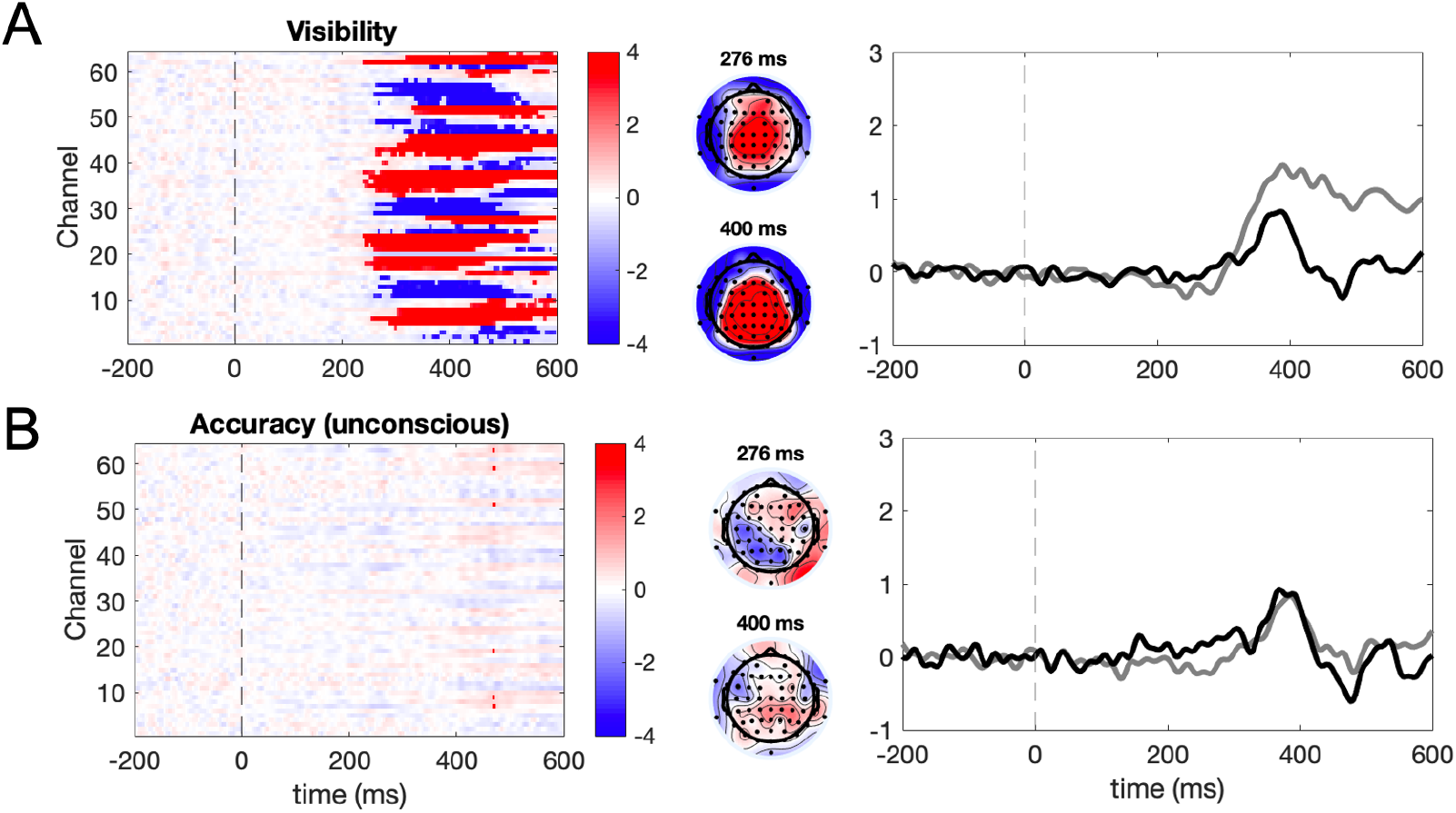
Event-related potentials. A) Single-trial visibility ratings (top panels) predicted ERPs 275 ms after stimulus onset. B) When only reportedly subliminal trials were included in the analysis (bottom panels), no strong effects were observed. The figures on the left show the results of the mass univariate mixed-effects regression models. Color denotes t value, and statistically significant effects (based on TFCE) are shown with a bright color. The middle panels show scalp topographies of the effects at 276 ms and 400 ms after stimulus onset (note that the electrode locations have been transposed: electrodes over the left scalp locations are contralateral to the stimulus). The rightmost panels show ERPs measured at the contralateral occipital cluster. The gray line depicts consciously seen (top panels) or correctly discriminated (bottom panels) trials, and the black line depicts not consciously seen (top panels)/incorrectly discriminated trials (bottom panels).

Similar pattern was revealed by event-related spectral perturbations (ERSP): Even though stimulus visibility clearly modulated ERSPs (Supplementary Figure 4), discrimination accuracy on subliminal trials did not statistically significantly modulate ERSPs at occipital electrodes (Supplementary Figure 5).

Next, we examined how prestimulus power influenced ERPs. If the prestimulus process that mediates accurate subliminal discrimination relates to fluctuations in cortical excitability, we should be able to observe that the contralateral occipital power is inversely related to ERP. Because the prestimulus EEG analyses indicated that broadband prestimulus offset correlated with subliminal discrimination, we first examined if this predicted ERPs (mixed-effects logit models with two predictors corresponding to offset in ipsilateral and contralateral occipital channels). These analyses did not yield any statistically significant effects.

We also examined whether single-trial oscillatory power in the same frequencies/time windows that predicted objective behavioral performance (ipsilateral electrode cluster at 1–3 Hz and contralateral electrode cluster at 8–12 Hz, 500–24 ms before stimulus onset) predicted ERP amplitude. We included only subliminal trials in the analysis. The results are shown in Figure 5. Contralateral occipital prestimulus power predicted ERP amplitude around 275 ms after stimulus onset in frontal electrodes. This effect is consistent with the observed relationship between ERPs and behavioral performance: As shown in Figure 4, conscious perception is associated with more positive ERPs in frontal sites after 275 ms. Figure 5 shows that the higher contralateral occipital prestimulus power, the weaker the ERP is frontal electrodes around 275 ms. Contralateral prestimulus power also statistically significantly modulated later responses (500 ms) in contralateral frontal electrodes. Ipsilateral occipital prestimulus power did not predict ERP amplitudes.

**Figure 5.**
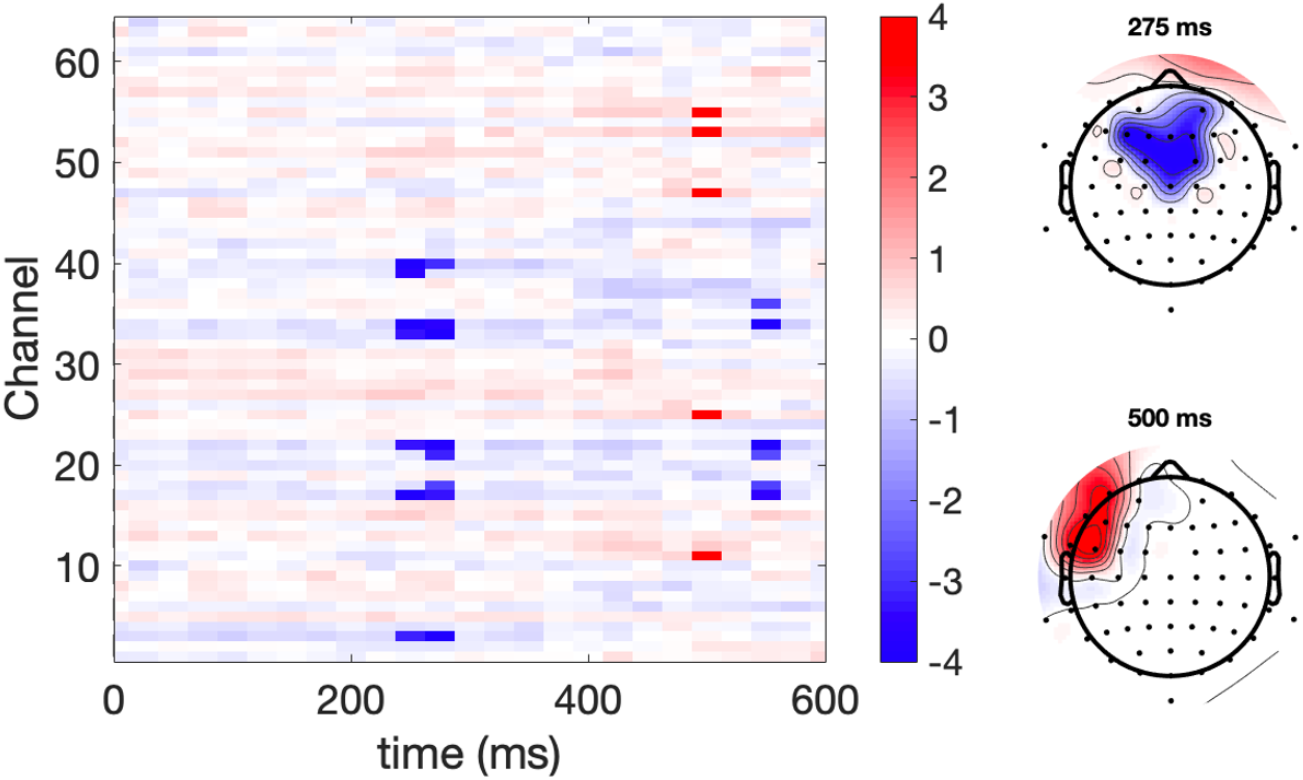
Influence of single-trial prestimulus oscillatory power on stimulus-evoked ERPs. Effect of contralateral occipital alpha prestimulus power on ERPs when the analysis includes only subliminal trials. The rectangular colormap plot shows the results of the mass univariate linear mixed-effects models. Color represents the t value. A strong color represents statistically significant effects, as determined by TFCE. Scalp maps show the distribution of the effects at specified time points (note that the electrode locations have been transposed: electrodes over the left hemisphere are contralateral to the stimulus).

## Discussion

We investigated what factors explain why behavioral visual discrimination accuracy can be on average above the level of chance, even though the participants subjectively report not seeing the stimulus (i.e. “subliminal perception”). Our results reveal the following: First, subliminal perceptual performance was not completely independent of conscious introspection, as measured by the signal detection theory. Second, lateralized neural activity before stimulus presentation, lateralization of stimuli, and previous trial’s discrimination accuracy predicted behavioral performance even when the participants reported just making guesses. Third, the accuracy of subliminal discrimination did not clearly modulate stimulus-evoked responses (ERP or ERPS), although as discussed below, this result may indicate lack of statistical power.

### Subliminal discrimination was not based on completely unconscious signals

Our behavioral results suggest that the performance on reportedly subliminal trials was not completely independent of the participants’ introspective abilities, at least based on the signal detection theoretic metrics of sensitivity and criterion (which were calculated based on subjective perceptual reports). This means that when the participants reported not seeing the stimuli, the above chance behavioral performance on these subliminal trials was likely nevertheless based on signals that the participants could introspect and base their response to. While it could be debated whether the nature of the information on what the participants based their decision on was visual or not (perhaps the experiences were type-2 like, as sometimes reported by blindsight patients), the key point is that the participants had some type of conscious access to information about the location of the stimulus on subliminal trials.

In general, signal detection theoretic framework implies that dissociations between subjective detection and forced-choice discrimination may be observed, not because there is a distinct unconscious perceptual capacity, but because the sensitivity of discrimination task is higher than the sensitivity of perceptual reports (Campion, Latto, & Smith, 1983; King & Dehaene, 2014; Phillips, 2018). Part of the difficulty in answering whether the signals on which the participants based their responses on subliminal trials were unconscious or not is that it requires us to artificially binarize a process that is in truth continuous. Our finding is in line with the observation that when a paradigm is used which minimizes the participants’ need to specify a criterion to label something as “conscious” or “unconscious”, no evidence for strictly unconscious perception is observed (Peters & Lau, 2015; Rajananda et al., 2020). Also, objective visual discrimination behavior and subjectively experienced perception evoked nearly identical ERP/ERSP responses, suggesting that the two are not strongly dissociated.

Our results are interesting when considered from the perspective of blindsight patients. Blindsight is typically assumed to demonstrate the existence of anatomical pathways that relay unconscious information to cortex, enabling the patients to use stimuli presented to their blind visual field to guide behavior (Ajina & Bridge, 2018; Ajina, Pestilli, Rokem, Kennard, & Bridge, 2015; Azzopardi & Cowey, 1997; Cowey, 2010; Danckert & Rossetti, 2005; Weiskrantz, Warrington, Sanders, & Marshall, 1974). Whether blindsight patients’ behavioral responding is based on completely unconscious percepts (Campion et al., 1983; Phillips, 2018), and whether similar mechanisms guide behavior in neurologically healthy individuals remains debated (Allen et al., 2020; Hurme, Koivisto, Henriksson, & Railo, 2020; Hurme, Koivisto, Revonsuo, & Railo, 2017, 2019; Lloyd, Abrahamyan, & Harris, 2013; for a review, see Railo & Hurme, 2021). An interesting open question is to what extent prestimulus processes also contribute to blindsight, and explain trial-by-trial variation in blindsight performance.

Finally, our results indicate that subliminal perception was more likely to be correct if the participant correctly discriminated the target in the previous trial. Furthermore, correct subliminal discrimination was more likely when stimuli were presented in the right hemifield. To what extent these factors reveal response biases, attentional or decision making biases, or long-term autocorrelations in the data (Schaworonkow et al., 2015) is unclear. Altogether, our results indicate that many different factors contribute to discrimination accuracy on reportedly subliminal trials. Subliminal discrimination accuracy could be predicted with AUC = 70% accuracy in the present study, which is very good performance given that we are essentially classifying trials which participants themselves label as “guesses”.

### Electrophysiological correlates

Baria et al. (2017) showed that participants’ behavioral responses on subliminal trials could be predicted from prestimulus activity patterns: when prestimulus activity resembled left stimulus-evoked responses, the participants were more likely to respond “left.” Because we transposed our electrode locations to correspond to contralateral versus ipsilateral space, our analysis was sensitive to the spatial pattern of the evoked response (although this also means that we may miss effects that are lateralized predominantly to left/right hemisphere). Our motivation to transpose electrode locations to contralateral versus ipsilateral space was largely that the lateralization of prestimulus power could explain how strongly the visual system responds to a visual stimulus in the contralateral visual field due to variation in cortical excitability. As discussed below, to what extent the present results are consistent with the “cortical excitability” explanation can be debated.

While previous research suggests that prestimulus alpha power modulates subjective detection and not objective discrimination (Iemi et al., 2017; Samaha et al., 2020), our results show that when the task is to discriminate lateralized targets, prestimulus alpha may also influence objective discrimination performance. In our study, we did not observe strong prestimulus effects for subjective visibility. However, as seen in Figure 2 and Supplementary Figure 2, subjective visibility was inversely associated with prestimulus alpha, although this effect did not quite reach statistical significance. Our results show that it is important to take into account the lateralization of prestimulus activity when examining prestimulus influences on perception when the task involves discrimination across hemifields. Although we observe that prestimulus activity predicts discrimination performance, the present results do not have to imply that prestimulus activity modulated perceptual *sensitivity* per se. Rather, because we transposed electrode locations, our analysis is sensitive to baseline excitability shifts (see, Iemi et al. 2017; Samaha et al., 2020) in both the contralateral and the ipsilateral hemispheres. While our results consistently indicate that contralateral occipital prestimulus activity was an important predictor of subliminal discrimination, also ipsilateral occipital activity may contribute to subliminal perception. Note that control analyses indicated that the prestimulus effects observed in the present study were possibly mediated by aperiodic broadband activity instead of oscillatory activity in the alpha band.

Early ERP waves (< 200 ms after stimulus onset) are assumed to reflect the addition of sensory-evoked neural activation on top of ongoing neural activity (Mäkinen, Tiitinen, & May, 2005; Shah et al., 2004). In addition to this, later ERP responses are modulated by asymmetric amplitude modulation of brain oscillations (Mazaheri & Jensen, 2008; Nikulin et al., 2007). Prestimulus neural activity may modulate ERPs through both of these mechanisms. Iemi et al. (2019) showed that while the power of prestimulus alpha- and beta-band activity influenced the earliest visual ERPs (C1 and N1 waves), the asymmetry of oscillations predominantly influenced later ERP waves.

We did not observe effects on early ERP amplitudes in the present study. One possibility is that our result is true, and there were not strong differences between early evoked activity in the correct vs. incorrect subliminal discrimination trials. This may suggest that correct responding in subliminal trials is determined by higher-level decision related processes rather than (relatively early) sensory processes. Interestingly, contralateral occipital prestimulus power predicted ERP amplitude in frontal areas when only the subliminal trials were included in the analysis. The topography and timing of this effect resembles correlates of conscious vision, and P3a ERP component, which has been interpreted as a correlate of orienting attention towards stimuli (Polich, 2007). This suggests that correct discrimination on subliminal trials may have to do with successfully attending the conscious “first-order” stimulus-related representations to enable correct discrimination decision (see, (Ko & Lau, 2012), for related mechanisms in blindsight patients). Prefrontal decision-making mechanisms can be activated by subliminal stimuli (Lau & Passingham, 2007; Van Gaal, Ridderinkhof, Scholte, & Lamme, 2010). Studies have also reported that activity in the frontal cortical areas correlates with subliminal behavioral performance (Lau & Passingham, 2006) and conscious vision (Persaud et al., 2011) in blindsight patients.

The influence of prestimulus oscillatory power on ERP amplitude could be a confound resulting from baseline correction of ERP’s: Trials with high prestimulus alpha power will also have non-zero mean amplitude due to the asymmetrical amplitude of the oscillations (Iemi et al., 2019). Therefore, it could be argued that when amplitude asymmetry is high (high alpha power), high amplitudes will be subtracted from the single-trial ERPs during baseline correction. Note that similar confound could also influence the ERP correlates of conscious and subliminal perception, either masking true effects or producing spurious findings.

It also remains possible that the strength of stimulus-evoked activity was modulated in the present study, but we were not able to detect this (likely very small) change due to e.g., limited statistical power of the mass univariate testing. In fact, Figure 4B suggests that correct subliminal perception was associated with amplified negative ERPs in contralateral occipital electrodes around 200 ms after stimulus onset. As noted, conscious perception of stimulus often correlates with ERPs in the same time-window (termed “VAN“; Förster et al., 2020), suggesting that correct subliminal discrimination is based on degraded conscious vision (Koivisto & Grassini, 2016). Similarly, while we did not observe statistically significant VAN in the present research, this may be due to lack of statistical power (due to mass-univariate testing) than lack of real amplitude difference in VAN time-window.

Because the aim of the present research was on subliminal perception, we restricted analysis to trials where participants reported not seeing the stimulus. An alternative method would have been to look for an interaction between, e.g., stimulus visibility and EEG measures (e.g., Benwell et al., 2017; Samaha et al., 2017). A demonstration of an interaction that reportedly visible and invisible trials are associated with different electrophysiological correlates could indicate that conscious and subliminal perception rely on clearly separate neural mechanisms. However, it is also possible that, e.g., the neural correlates of above-threshold perception and near-threshold perception differ, not because these are truly different forms of perception, but simply because they are two different perceptual decisions based on different perceptual contents (e.g., one based on clear perception of a stimulus, and other based severely degraded percept).

### Conclusion

We have shown that neural activity before stimulus presentation can be used to predict whether an individual will correctly discriminate the location of a stimulus they report not seeing. Furthermore, our results indicate that performance on these reportedly unconscious trials may actually depend on processes that a participant can consciously introspect. This way, our finding speaks against the popular notion of a separate, unconscious perceptual capacity, and emphasizes that visual behavior forms a continuum that is tightly linked with the associated subjective feeling of seeing.

## Acknowledgments

The study was funded by the Academy of Finland (grant #308533). The authors also thank two reviewers for very helpful comments.

## Supplementary information

**Supplementary Figure 1.**
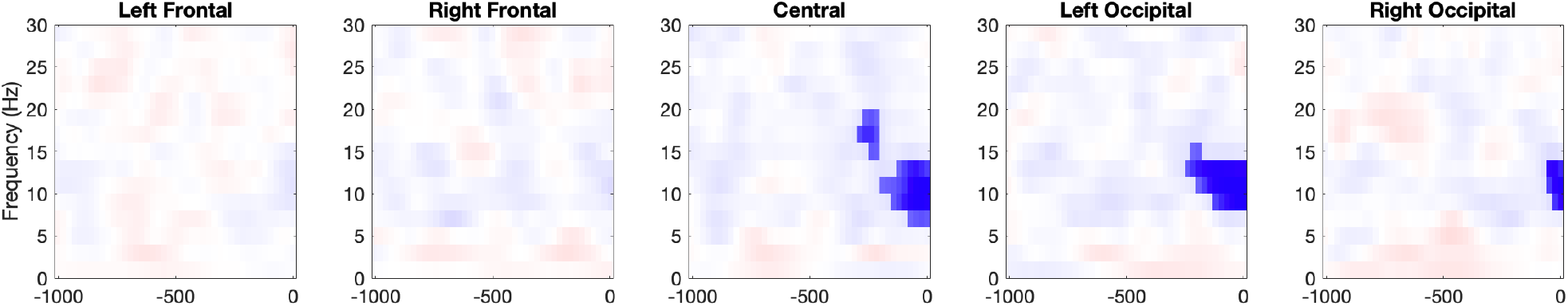
Predicting the accuracy of objective discrimination (on subliminal trials) based on prestimulus power. This analysis was performed on non-transposed electrode locations. Color represents the t value. A bright color represents statistically significant effects according to TFCE. Color scale is the same as in the main article (i.e., color limits are [−4 4]).

**Supplementary Figure 2.**
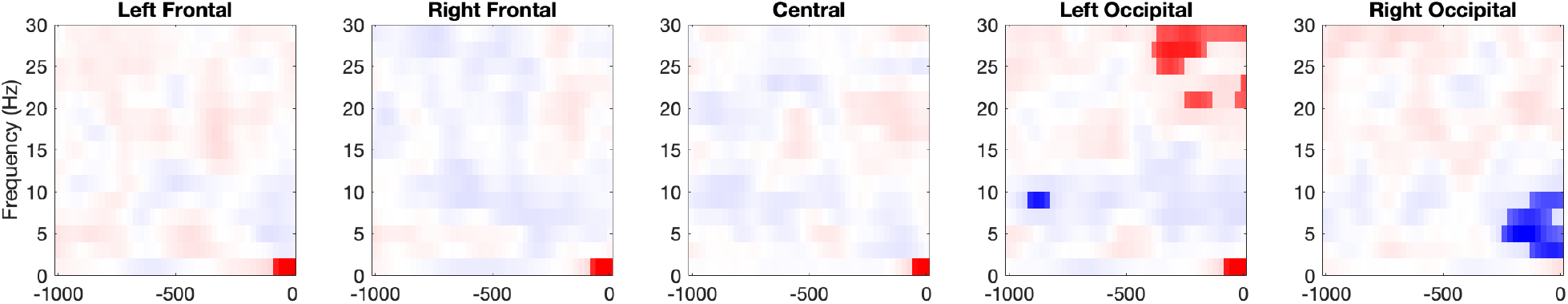
Predicting (binarized) conscious stimulus visibility based on prestimulus power. This analysis was performed on non-transposed electrode locations. Color represents the t value. A bright color represents statistically significant effects according to TFCE. Color scale is the same as in the main article (i.e., color limits are [−4 4]).

**Supplementary Figure 3.**
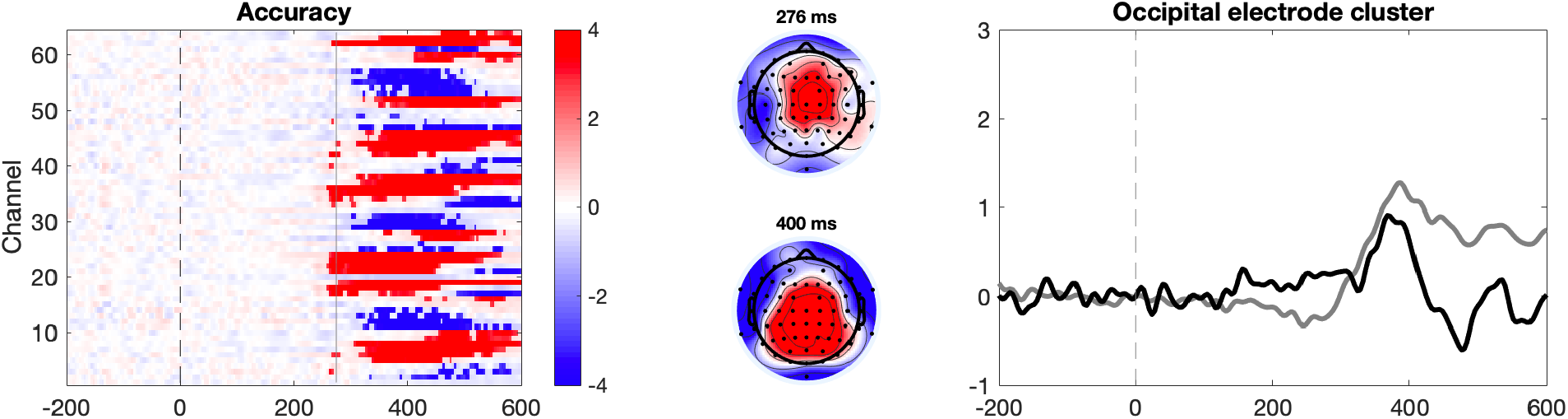
How event-related potentials were modulated by discrimination accuracy (analysis include both seen and not-seen trials). Bright color represents statistically significant effects according to TFCE. In the right panel black line is incorrect response, and gray line is correct response.

**Supplementary Figure 4.**
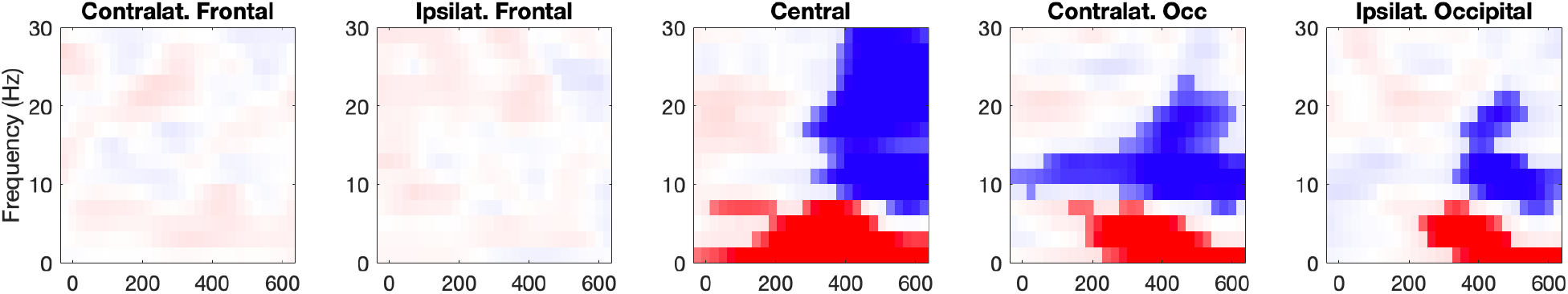
How ERSPs were modulated by binarized subjective visibility. Bright color represents statistically significant effects according to TFCE. Color scale is the same as in the main article (i.e., color limits are [−4 4]).

**Supplementary Figure 5.**
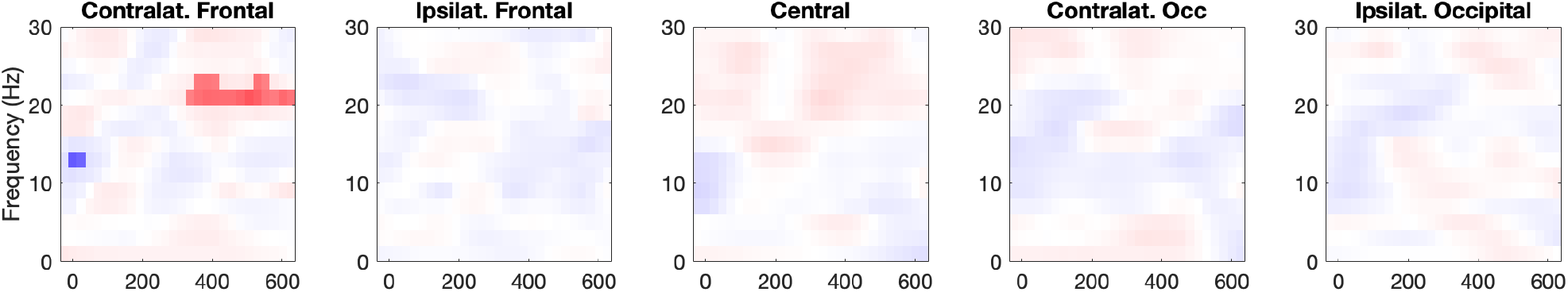
How ERSPs were modulated by discrimination accuracy on subliminal trials. Bright color represents statistically significant effects according to TFCE. Color scale is the same as in the main article (i.e., color limits are [−4 4]).

